# Accelerated FRET-PAINT Microscopy

**DOI:** 10.1101/376913

**Authors:** Jongjin Lee, Sangjun Park, Sungchul Hohng

## Abstract

Recent development of FRET-PAINT microscopy significantly improved the imaging speed of DNA-PAINT, the previously reported super-resolution fluorescence microscopy with no photobleaching problem. Here we try to achieve the ultimate speed limit of FRET-PAINT by optimizing the camera speed, dissociation rate of DNA probes, and bleed-through of the donor signal to the acceptor channel, and further increase the imaging speed of FRET-PAINT by 8-fold. Super-resolution imaging of COS-7 microtubules shows that high-quality 40-nm resolution images can be obtained in just tens of seconds.

---

Different types of super-resolution fluorescence microscopy techniques have been developed to overcome the diffraction limit of optical microscopy.^1-7^ The achievement, however, was obtained by sacrificing imaging speed and total observation time; with increased optical resolution, the imaging speed is generally slowed-down and the photobleaching problem of fluorophores becomes exacerbated resulting in the limited total imaging time. DNA-PAINT (Point Accumulation for Imaging in Nanoscale Topography^8^) technique has overcome the photobleaching problem by using transient binding of a fluorescently labeled short DNA strand (imager strand) to a docking DNA strand conjugated to target molecules.^9^ The binding rates of DNA probes, however, are notoriously slow, and as a result, DNA-PAINT has an extremely slow imaging speed (1-3 frames per hour), impeding widespread usage of DNA-PAINT in biological imaging. To solve this problem of DNA-PAINT, FRET-PAINT microscopy has been introduced independently by two groups.^10,11^ In this technique, two short DNA strands labeled with donor and acceptor are used as fluorescence probes. Because only the acceptor signal is used for single-molecule localization, more concentrated DNA probes could be used, resulting in a 30-fold increase in imaging speed compared to DNA-PAINT.^10^

The ultimate speed limit of FRET-PAINT has not been characterized yet. The imaging speed of FRET-PAINT is influenced by the camera speed, dissociation rate of DNA probes, and maximum concentration of DNA probes. In this paper, we optimize the three factors to reach the speed limit of FRET-PAINT imaging, and as a result, report a super-resolution fluorescence microscopy that can provide 40-nm resolution images in tens of seconds. In this process, we recognized the previously uncharacterized photo-induced damage of DNA probes, which currently limits both the imaging speed and the observation time of FRET-PAINT.

The experimental scheme of FRET-PAINT and instrumental setup are briefly presented in Figure 1a. In the previous work, we used an EMCCD (iXon Ultra DU-897U-CS0-#BV, Andor) with a maximum frame rate of 56 Hz and 512 x 512 imaging area. Due to slow dissociation of DNA probes, however, actual frame rate used was 10 Hz. In this work, we replaced the EMCCD with an sCMOS camera (ORCA-Flash 4.0 V2, Hamamatsu) with a maximum frame rate of 400 Hz for the same size of an imaging area. Due to the photo-induced damage of DNA probes that will be explained later in more detail, however, the maximum frame rate used was 200 Hz. To compensate for short exposure time, illumination intensity should be increased proportionally to frame rate. For the same reason of photo-induced DNA damage, we used an illumination power of 1.5 kW/cm^2^, just 3.3-fold increase from 460 W/cm^2^ that was used in the previous work.

**Figure 1.**
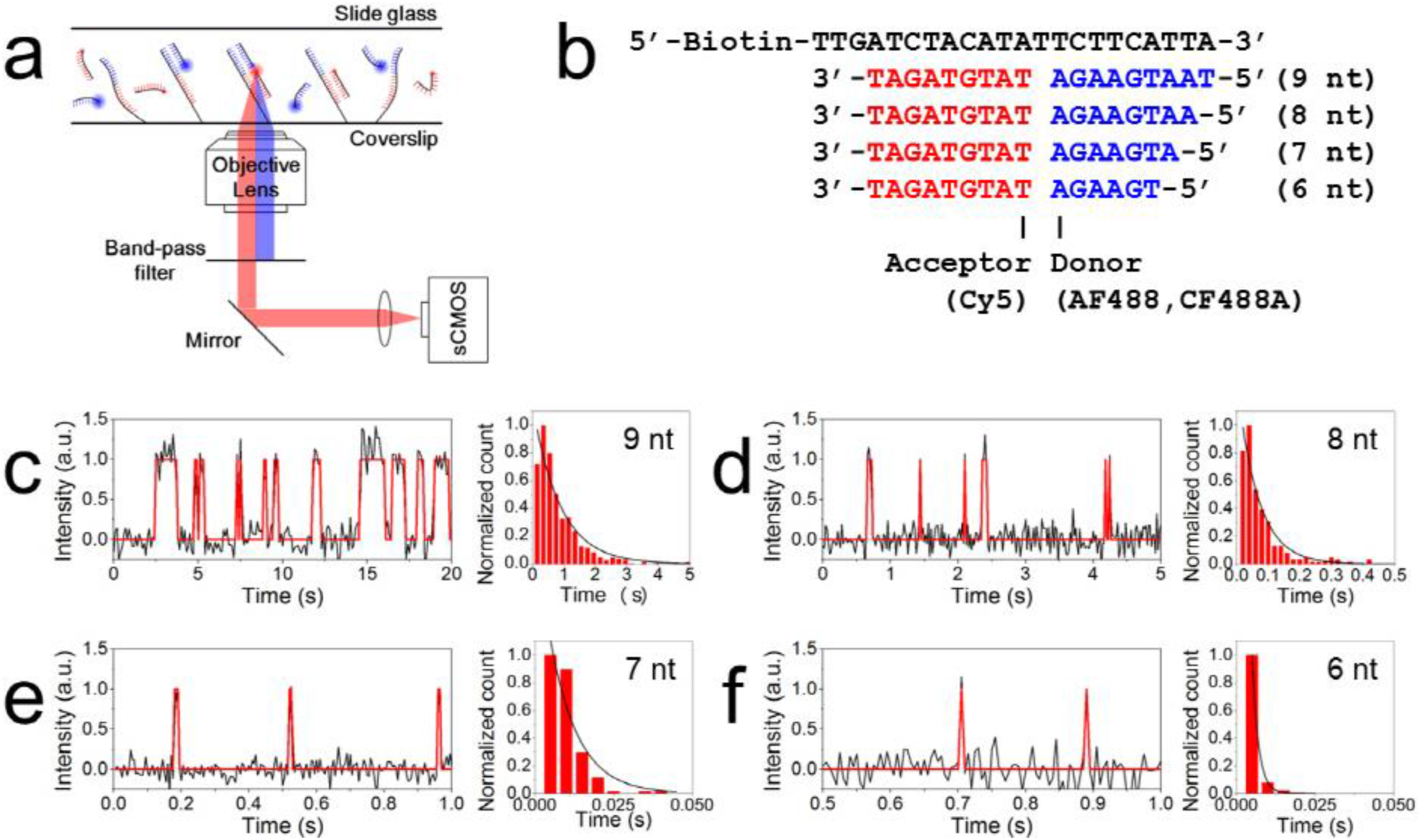
Accelerated dissociation of donor strands. (a) A scheme of FRET-PAINT microscopy. Acceptor fluoresces only via FRET and its signal is collected by a high-speed sCMOS camera. Donor signal is rejected by a band-pass filter. (b) DNA strands used for the experiments: docking (black), donor (blue), and acceptor (red) strands. A length of donor strand was controlled by truncating the 5’-end of the donor strand. Acceptor and donor fluorophores are labeled at the designated positions. (c-f) Dissociation time of donor strands with the length of 9 nt (c), 8 nt (d), 7 nt (e), and 6 nt (f). Left panels show representative FRET time traces, in which high and low FRET states correspond to the bound and unbound states, respectively. Right panels show histograms of dissociation times. The dissociation times were obtained by fitting the histograms with an exponential decay function: 800 ms (9 nt), 66 ms (8 nt), 8.9 ms (7 nt), and 2 ms (6 nt).

To fully utilize the increased frame rate of an sCMOS camera, the switching rate of DNA probes should be increased as well; if the dissociation of DNA probe is slow, single-molecule spots start to overlap at lower probe concentrations, limiting the overall imaging speed. We determined the dissociation times of donor strands with various lengths. Four different donor strands were tested (Figure 1b, blue). Figure 1c-f show representative time traces (left) and histograms of dissociation time of the donor strands (right). The dissociation times obtained were 800 ms (9 nt), 66 ms (8 nt), 8.9 ms (7 nt), and 2.0 ms (6 nt). We selected 7 nt donor strands for the frame rate of 100 or 200 Hz used in this work.

Background noise coming from floating donor and acceptor strands limits the maximum probe concentration that can be used. To reduce the background noise, and as a result to increase the maximum probe concentrations to give reasonable signal-to-noise ratio (SNR), we first tried different donor-acceptor pairs other than the Alexa Fluor 488 (AF488, Invitrogen)-Cy5 (GE Healthcare) pair used in the previous work. In terms of background noise, the more the spectral separation of donor and acceptor emissions, the better SNR. Through paper research, CF488A (Biotium) and CF660R (Biotium) were selected as candidates to replace AF488 and Cy5, respectively. Figure 2a compares excitation (dashed lines) and emission (solid lines) spectra of AF488 (black), CF488A (red), Cy5 (magenta), and CF660R (violet). The absorption and emission spectra of CF488A is blue-shifted to those of AF488 whereas their extinction coefficients are similar at the peaks. On the other hand, the emission spectrum of CF660R is red-shifted to that of Cy5. As an additional effort to improve SNR, we also replaced a 640 nm long-pass filter (green dashed line) with a 700/75 band-pass filter (green solid line). Since the band-pass filter has a red-shifted cut-on wavelength than the long-pass filter, some portion of acceptor signal is lost by the replacement, but we expected the reduction of donor bleed-through would increase SNR at high donor strand concentrations. As expected from the fact that CF488A has larger extinction coefficient than AF488 at 473 nm, the CF488A-Cy5 pair gave more photons than the AF488-Cy5 pair at the same excitation power (Figure 2b). The background noise, and thus the signal to noise ratio, were also improved with the CF488A-Cy5 pair than the AF488-Cy5 pair (Figure 2c,d). It is noticeable that the optimization process mentioned above removed the donor bleed-through almost completely, and as a result, the dependence of SNR on donor concentration was very weak (Figure 2d). Contrary to our expectation, we found that replacement of Cy5 with CF660R did not improve SNR because CF660R has higher direct excitation than Cy5 at 473 nm (Figure S1). Since CF660R has lower direct excitation than Cy5 at 488 nm, we expect that CF660R may provide better performance if we use a 488-nm excitation laser instead of the 473-nm laser in a future work. In this work, we exclusively used the CF488A-Cy5 pair at 473 nm excitation.

**Figure 2.**
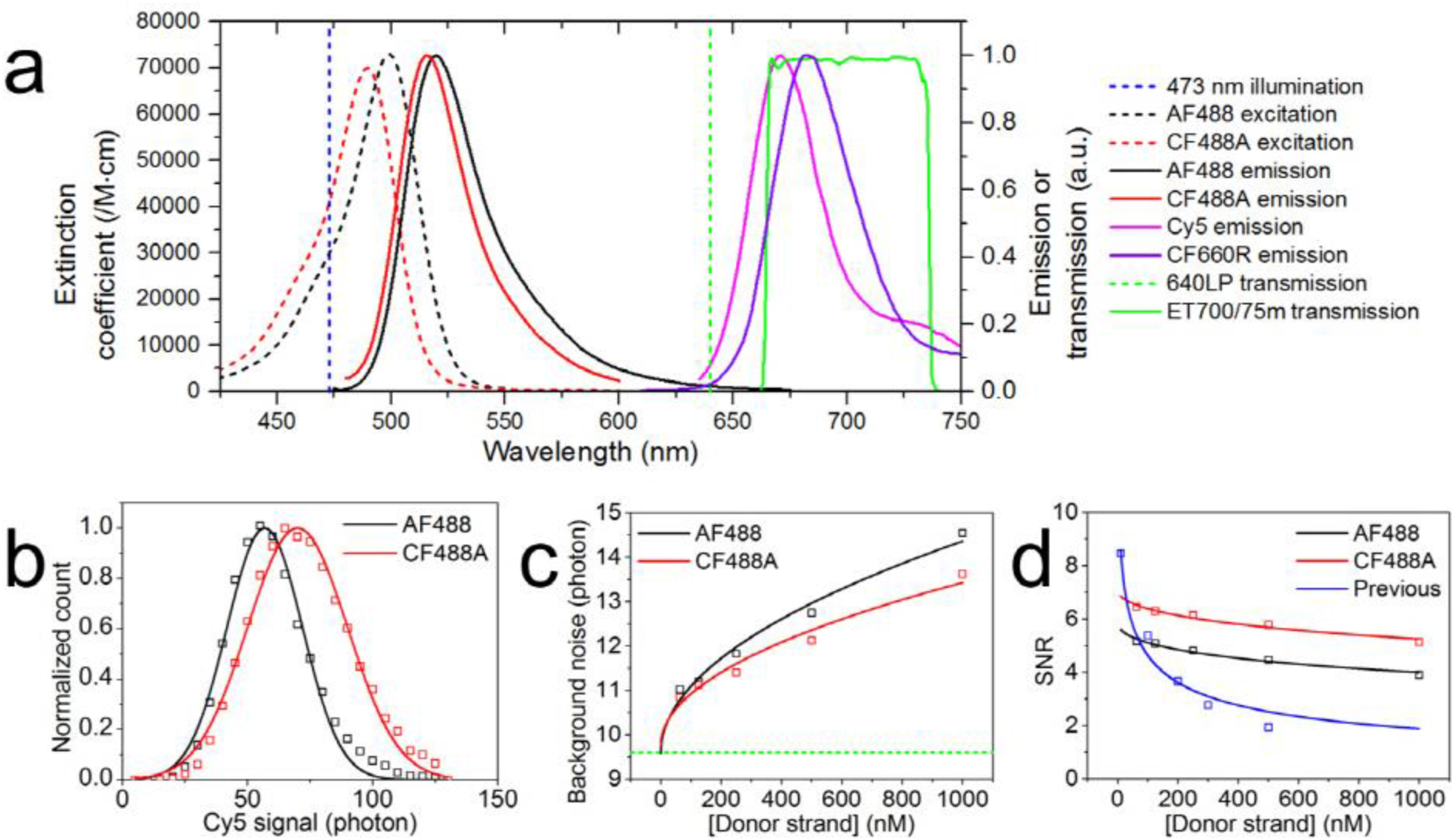
Improved signal-to-noise ratio (SNR). (a) Excitation (dashed lines) and emission spectra (solid lines) of donor (AF488, black; CF488A, red) and acceptor (Cy5, magenta; CF660R, violet) fluorophores. The vertical blue dashed line indicates 473 nm excitation wavelength, the vertical green dashed line indicates cut-on wavelength of a 640 nm long-pass filter, and the green solid line indicates the transmission curve of a 700/75m band-pass filter. (b) Acceptor signal of the AF488-Cy5 (black) and CF488A-Cy5 (red) pairs at 1.5 kW/cm^2^ excitation power recorded with an sCMOS camera and a band-pass filter. The signal is defined as the amplitude of a 2D Gaussian function of each single-molecule spot. Open squares indicate measured values and solid lines indicate fitted curves with Gaussian function. The CF488A-Cy5 pair yields the higher intensity. (c) Background noise of the AF488-Cy5 (black) and CF488A-Cy5 (red) pairs with an sCMOS camera and a band-pass filter. The background noise is defined as the FWHM of a Gaussian function of the background signal. Open squares indicate measured values and solid lines indicate fitted curves with a square root of donor strand concentration. CF488A-Cy5 pair yields lower background noise. Horizontal green dashed line indicates background noise without donor and acceptor strands, which is mainly caused by autofluorescence coming from a coverslip. (d) SNR of the AF488-Cy5 (black) and CF488A-Cy5 (red) pairs recorded with an sCMOS camera and a band-pass filter and that of the AF488-Cy5 pair (blue) recorded with an EMCCD camera and a long-pass filter. SNR is defined as the ratio of the signal to the background noise. Open squares indicate calculated values and solid lines indicate fitted curves with an inverse square root function of donor strand concentration. The CF488A-Cy5 pair with an sCMOS camera and a band-pass filter yields the highest SNR at high donor strand concentration.

To characterize the improved imaging speed of the new microscope, we compared the imaging speed of the new FRET-PAINT microscope with the previous one. As a model system, microtubules of COS-7 cells were imaged. 7 nt donor strands were used for the new microscope whereas 9 nt donor strand was used for the old one. Figure 3a shows a super-resolution image obtained with the old microscope using a 10 Hz frame rate and 1 minute acquisition time. For the imaging, 30 nM AF488-labeled donor strands and 20 nM Cy5-labeled acceptor strands were used. Figure 3b and Figure 3c show super-resolution images obtained with the new microscope using 100 Hz frame rate for Figure 3b or 200 Hz for Figure 3c. For the imaging, the total data acquisition time was 1 minute, and 300 nM CF488A-labeled donor strands and 300 nM Cy5-labeled acceptor strands were used. As clear from the Figures, the new microscope provided higher quality images than the previous FRET-PAINT setup in a shorter time. To show the improved image qualities in more detail, time-lapse images of the boxed regions of Figure 3a-c are also shown in Figure 3d-f, respectively. To quantitatively compare image qualities of Figure 3a-c, we compared the image resolutions as a function of image acquisition time (Figure 3g). The resolutions were obtained using the Fourier ring correlation method.^12,13^ It is noticeable that the resolution arrived at the limit (42 nm for 100 Hz, 46 nm for 200 Hz) after 20-30 s with the new FRET-PAINT setup whereas the resolution still decreases even after 60 s with the previous FRET-PAINT setup. In principle, the resolution defined by Fourier ring correlation method is affected by both the localization precision and the localization density.^12-15^ The localization density is linearly proportional to the imaging time (Figure 3h) whereas the localization precision is time-independent. Therefore we can conclude that for tens of seconds imaging time the image resolution is determined by the localization precision in a new microscope. For the same image acquisition time, on the other hand, the image resolution is determined by the localization density in the old microscope. Localization density as a function of imaging time in Figure 3h provides another way to compare the imaging speed of the microscopes. The localization rate was increased by 5.4 times for the 100 Hz imaging, and 8 times for the 200 Hz imaging.

**Figure 3.**
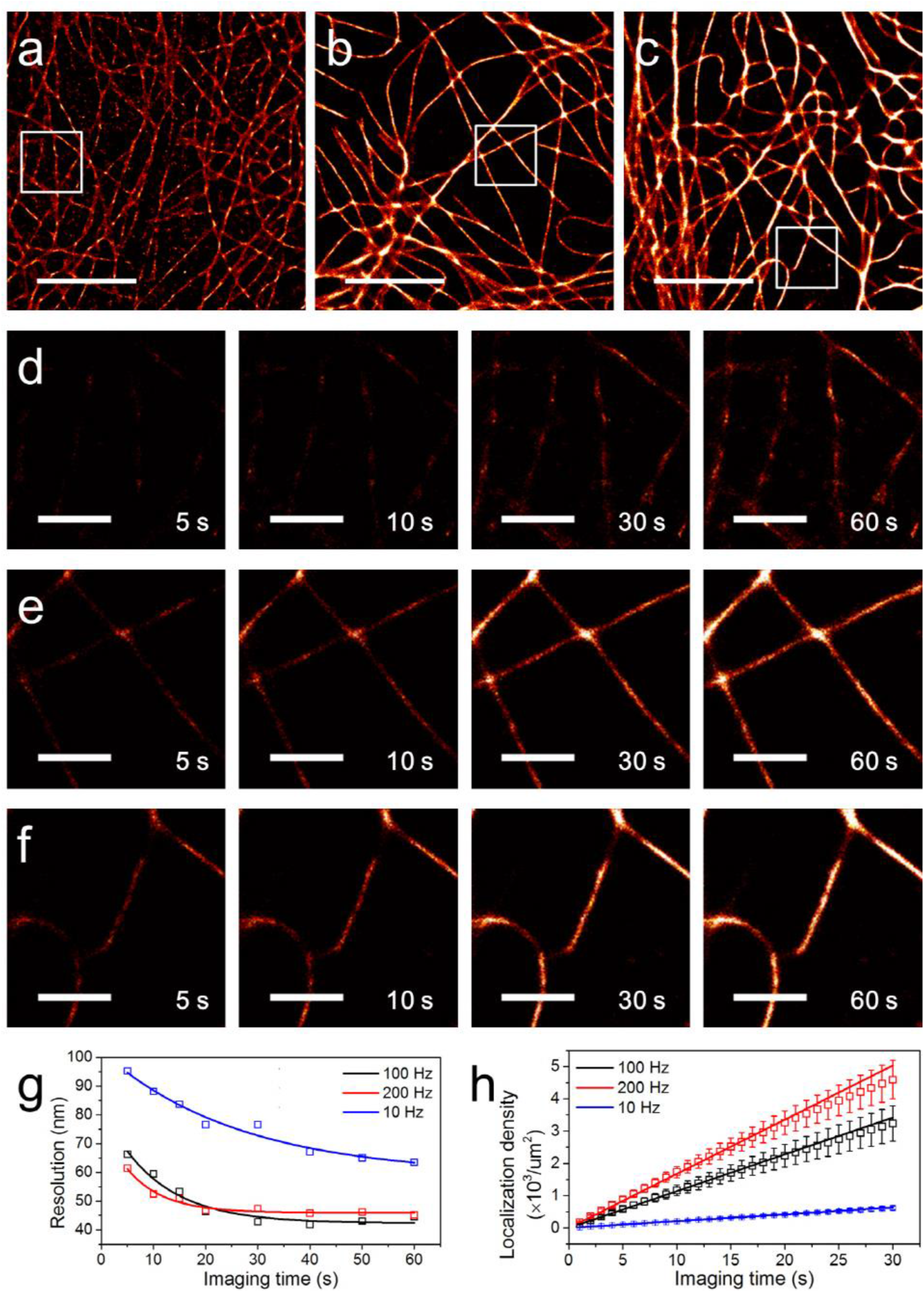
Characterization of the imaging speed of a new microscope. Super-resolution microtubule images of fixed COS-7 cells were used as a model system. (a) The image was reconstructed from 600 frames recorded at a frame rate of 10 Hz with a previous microscope (an EMCCD camera, a long-pass filter, 30 nM 9 nt AF488 donor strands, 20 nM 10 nt Cy5 acceptor strands). (b,c) The images were reconstructed from 6000 frames recorded at a frame rate of 100 Hz (b) or 12000 frames recorded at a frame rate of 200 Hz (c) with a new microscope (an sCMOS camera, a band-pass filter, 300 nM 7 nt CF488A donor strands, 300 nM 9 nt Cy5 acceptor strands). All images were reconstructed using ThunderSTORM^18^ with maximum likelihood fitting method. Total imaging time is 60 s for all images. (d-f) Time-lapse images of the boxed regions in panels (a-c) at the specified imaging time. (g) Image resolutions of panels (a-c) using Fourier ring correlation method as a function of the imaging time. Open squares indicate measured value and solid lines indicate fitted curves with an exponential decay function. (h) A localization density as a function of the imaging time (100 Hz, black; 200 Hz, red; 10 Hz, blue). The localization density is defined as the number of localization events per um^2^. To minimize the influence of the region of interest selected for data analysis, the localization density was calculated from 10 different regions of 5 different cells. Squared boxes indicate the average and error bars indicate the standard deviation. The increase rates of the localization density were 21 (10 Hz), 114 (100 Hz), and 168 (200 Hz) localizations/um^2^/s. We obtained 5.4 times increase for 100 Hz imaging, and 8 times increase for 200 Hz imaging compared to the old microscope. Scale bars: 5 um (a-c), 1 um (d-f).

In summary, we developed a high-speed FRET-PAINT microscope that can provide localization-accuracy limited super-resolution images in tens of seconds. For the achievement, we optimized several experimental parameters such as the camera speed, dissociation time of donor strands, and bleed-through of donor signals to the acceptor channel. Then, have we arrived at the ultimate speed limit of FRET-PAINT microscopy? Figure 2d shows that we could use much higher donor strand concentrations than 300 nM without compromising SNR. The localization accuracy could be also improved by collecting more photons. By using 6 nt donor stand, the donor strand switching rate could be also increased. Incorporation of all these change into the microscope to increase the imaging speed, however, requires higher excitation intensity to compensate for the decreased photon number caused by the reduced binding lifetime of the probes. Unfortunately, we found that this simple scheme did not work; DNA probes used in FRET-PAINT were damaged by the high-intensity excitation laser (Figure S2a). Since the injection of fresh DNA probes did not solve the problem (Figure S2b) and the background noise of fluorophores did not decrease (Figure S2c), the damage is not simple photobleaching of fluorophores but seems to be the loss of base-paring capability of the docking strand. Interestingly, we found that the photo-induced damage exhibited sample-to-sample variation (Figure S2d). Finding of a way to systematically solve the photo-induced problem will enable us to realize sub-millisecond image acquisition for super-resolution imaging. When combined with a recently-developed real-time confocal microscopy,^16,17^ our accelerated FRET-PAINT microscopy will provide a way to reconstruct three-dimensional structures of thick neural tissue samples with both high speed and high resolution.

## Supporting Information

The Supporting Information is available free of charge.

Experimental details and supporting figures (PDF)

## ACKNOWLEDGMENT

This work was supported by a Creative Research Initiative grant (Physical Genetics Laboratory, 2009-0081562) to S.H.

---

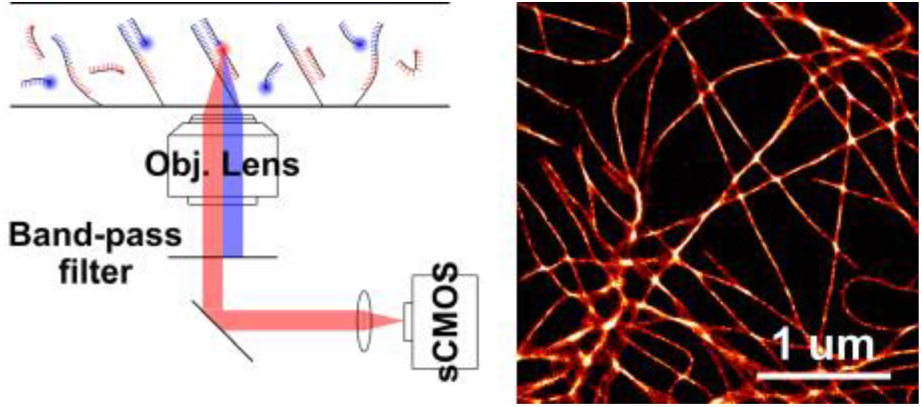

**Author Contributions**
J.L. and S.H. designed experiments. J.L. and S.P. performed experiments and analyzed data. S.H. supervised the research. J.L. and S.H. wrote the paper.

